## Supplemental Information for "Intrinsic dynamic shapes responses to external stimulation in the human brain"

### Supplementary data

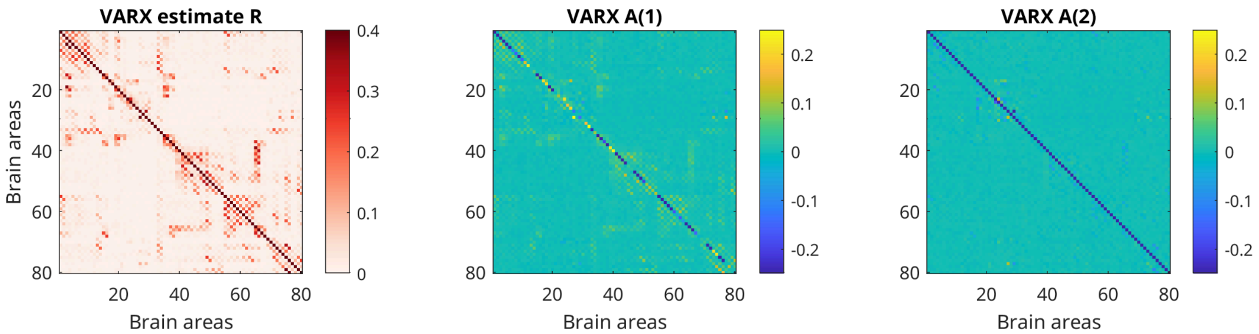


**Figure S1: Connectivity of stimulated neural mass model:** On the left is the same effect matrix $R$ as in Fig. 2B, except now the diagonal element is shown. The two matrices on the center and right indicate the $\mathbf{A}$ filter matrix for delays 1 and 2. The color axis has been limited to ±0.25 for visibility of off-diagonal elements.

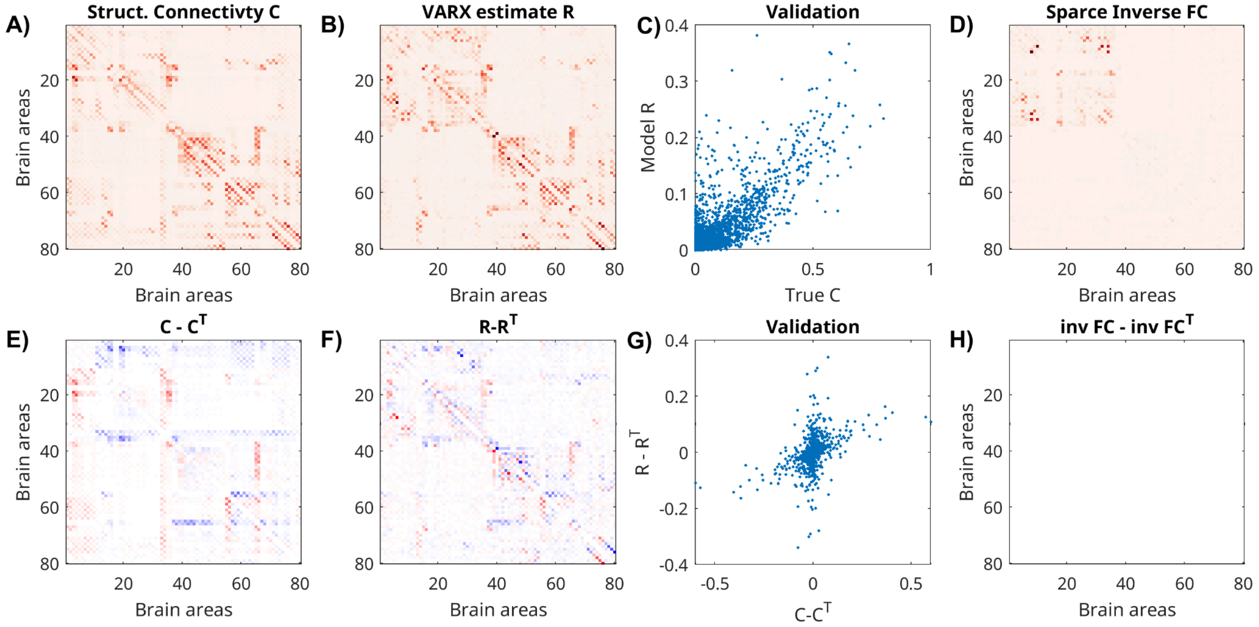


**Figure S2: Connectivity of stimulated neural mass model with asymmetric structural connectivity.** A) Same as in Fig. 2A, however, rows 1:10, 65:66, 33:36, 55:56 have been downscaled by a factor of 0.1 to enhance the asymmetry of the structural connectivity matrix for nodes that had larger connectivity between distant brain areas. Note again that the diagonal in $R$ is omitted as it is also missing in the structural connectivity matrix of the simulation. B) (C) Spearman correlation drops to r=0.54 (from 0.69 in Fig. 2). E) Asymmetry is highlighted by subtracting the transpose (same as in Fig. 7). F) The VARX largely recovers the sign of the asymmetry. G) But a number of nodes are misestimated. H) the sparse inverse covariance by definition is symmetric and does not capture any asymmetry.

| Patient ID | Age | Sex | Total Length of Recordings [min] | Number of Channels |
|---|---|---|---|---|
| Pat_1 | 58 | M | 58.6 | 153 |
| Pat_2 | 22 | M | 48.6 | 164 |
| Pat_5 | 48 | M | 58.6 | 334 |
| Pat_6 | 36 | F | 58.6 | 189 |
| Pat_7 | 43 | M | 58.6 | 132 |
| Pat_8 | 41 | F | 64.7 | 154 |
| Pat_9 | 50 | M | 58.6 | 267 |
| Pat_9_02 | 51 | M | 58.6 | 271 |
| Pat_10 | 24 | M | 53.6 | 192 |
| Pat_11 | 37 | M | 58.6 | 192 |
| Pat_11_02 | 37 | M | 58.6 | 111 |
| Pat_12 | 52 | F | 48.6 | 198 |
| Pat_13_02 | 24 | M | 58.6 | 296 |
| Pat_14 | 20 | M | 58.6 | 207 |
| Pat_15 | 56 | M | 53.6 | 100 |
| Pat_16 | 43 | F | 39.3 | 261 |
| Pat_17 | 27 | F | 58.6 | 220 |
| Pat_18 | 28 | F | 58.6 | 227 |
| Pat_18_02 | 30 | F | 58.6 | 71 |
| Pat_19 | 46 | M | 48.6 | 230 |
| Pat_20 | 35 | F | 54.3 | 231 |
| Pat_21 | 48 | M | 58.6 | 323 |
| Pat_22 | 19 | M | 48.6 | 75 |
| Pat_23 | 36 | F | 54.4 | 175 |
| Pat_23_03 | 36 | F | 48.7 | 260 |
| Pat_24 | 59 | F | 39.3 | 60 |

**Table S1. Demographics, length of recordings and number of recording channels.** Data from patients 9, 11, 18 and 23 were recorded from two reimplants each at different times.

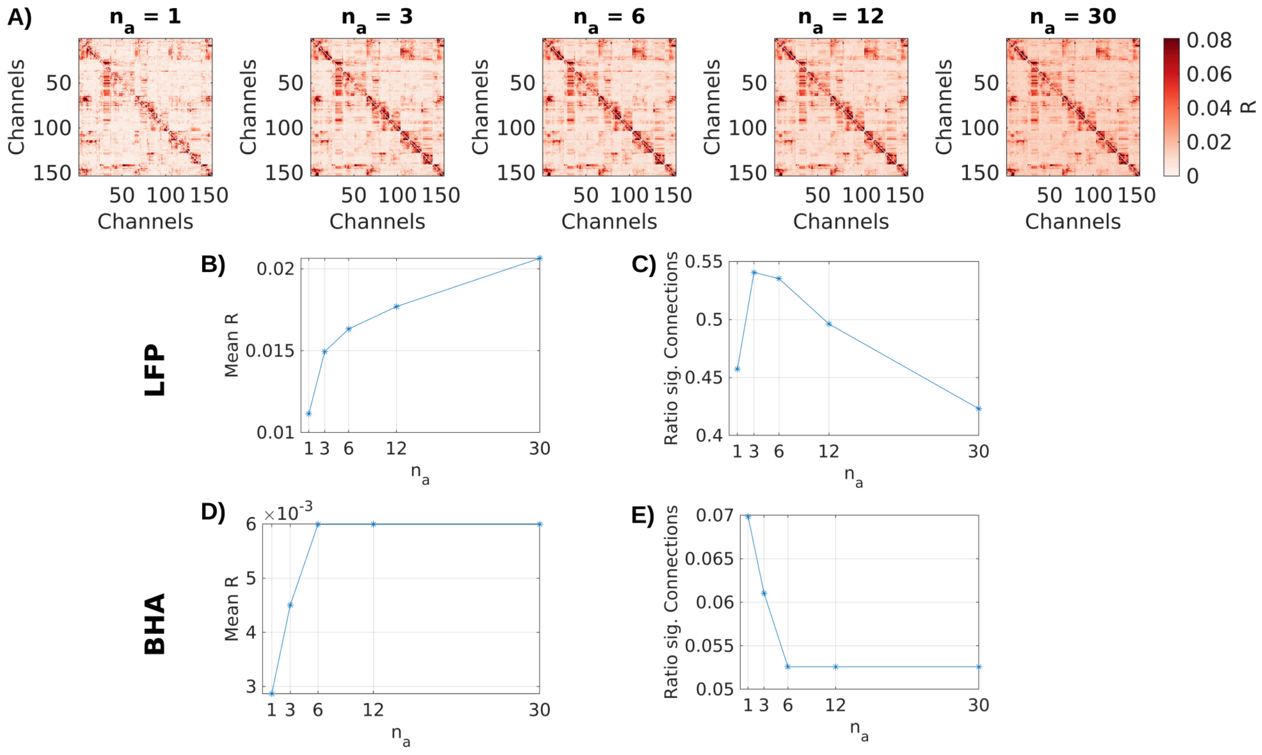


**Figure S3: Choice of $n_a$.** A) Effect size R of connections in LFP data for an example patient using different values for the delays $n_a$. B) Mean effect size R, and C) ratio of significant ($p<0.001$) channels across all channels for different $n_a$ for LFP data. D) and E) in BHA.

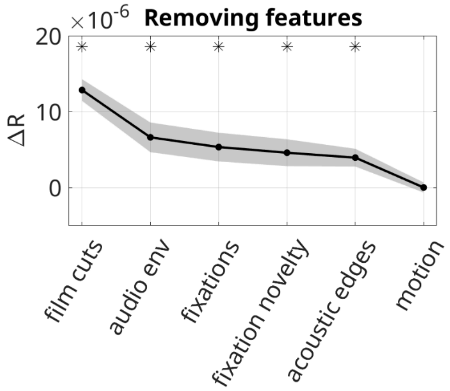


**Figure S4: Effect of individual extrinsic features on effect size of recurrent connections.** Removing regressors for film cuts ($\Delta R=-12.8*10^{-6}$, $p<0.0001$, N=26), the auditory envelope ($\Delta R=-6.63*10^{-6}$, $p=0.0002$, N=26), acoustic edges ($\Delta R=-3.95*10^{-6}$, $p=0.007$, N=26), fixation novelty ($\Delta R=-4.6*10^{-6}$, $p=0.005$, N=26), or fixation onset ($\Delta R=-5.6*10^{-6}$, $p=0.0009$, N=26) significantly increases the effect size compared to the VARX model including all features. Removing the motion regressor does not show this effect ($\Delta R=-0.02*10^{-6}$, $p=0.64$, N=26). FDR correction, $\alpha=0.05$. Black line shows the mean increase of effect size ΔR. Gray shaded area shows the standard error of the mean of ΔR across 26 patients. Stars indicate significant changes in effect size.

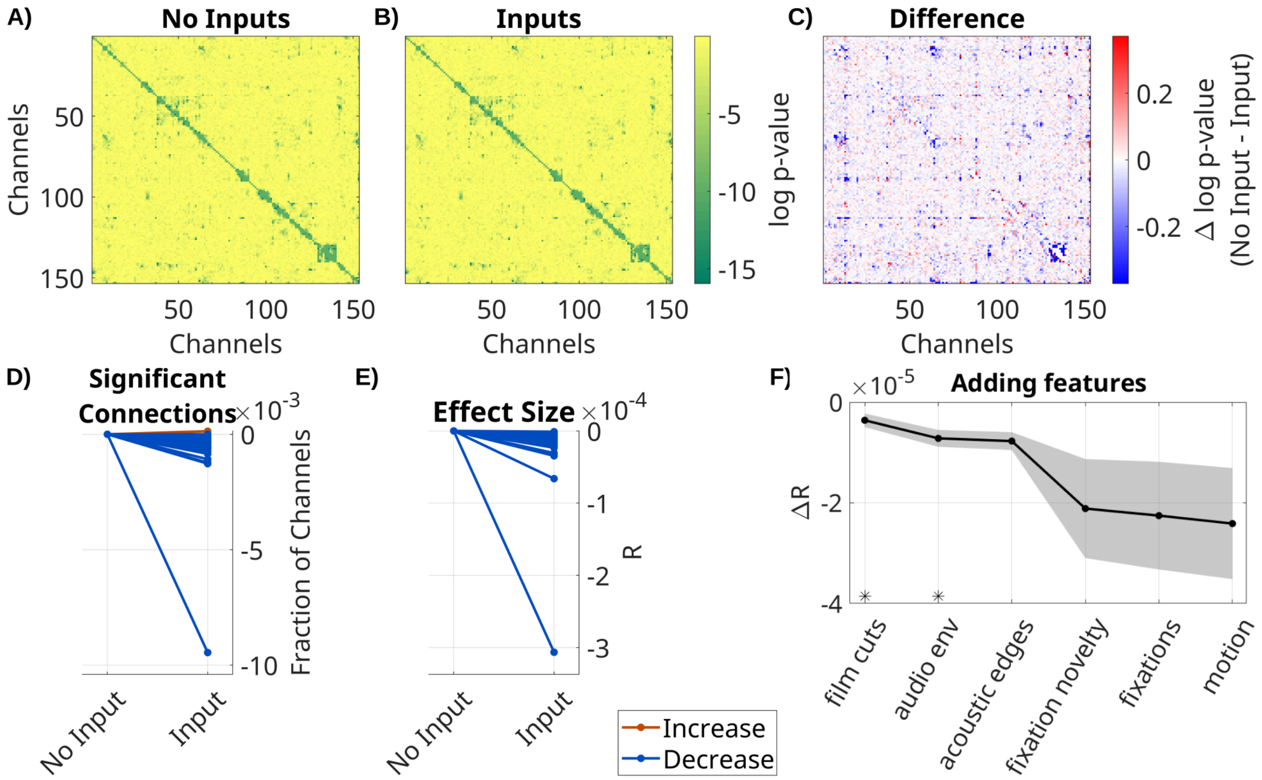


**Figure S5: Spurious recurrent BHA connectivity in $\mathbf{A}$ is accounted for when modeling the effect of input with $\mathbf{B}$.** Same analysis as in Figure 3 with BHA data. A) $p$-values for each connection in $\mathbf{A}$ for VARX model without inputs on one patient (Pat_1); B) for VARX model with inputs; C) difference. Both models are fit to the same data. D) Fraction of significant recurrent connections in VARX models with and without inputs (difference: median=$-3.7*10^{-4}$, p<0.0001, N=26, Wilcoxon). E) Effect size $R$ over all electrodes between VARX models with and without inputs (difference: median=$-1*10^{-5}$, p<0.0001, N=26, Wilcoxon). Each line is a patient, with color indicating an increase or decrease. Values in D) and E) have been normalized to models without input. F) Difference between the VARX model without input and VARX models successively adding inputs. Black line shows mean across patients, shaded gray area the standard error of the mean. Stars indicate features that further reduce effect size over the previously added feature with statistical significance (Wilcoxon rank sum test p<0.05). Negative values indicate a decrease in connectivity strength when the extrinsic inputs are accounted for. Adding film cuts ($\Delta R=-3.6*10^{-6}$, p<0.0001, N=26, FDR correction, α=0.05), and the auditory envelope ($\Delta R=-3.6*10^{-6}$, p=0.002, N=26, FDR correction, α=0.05) significantly decrease R values.

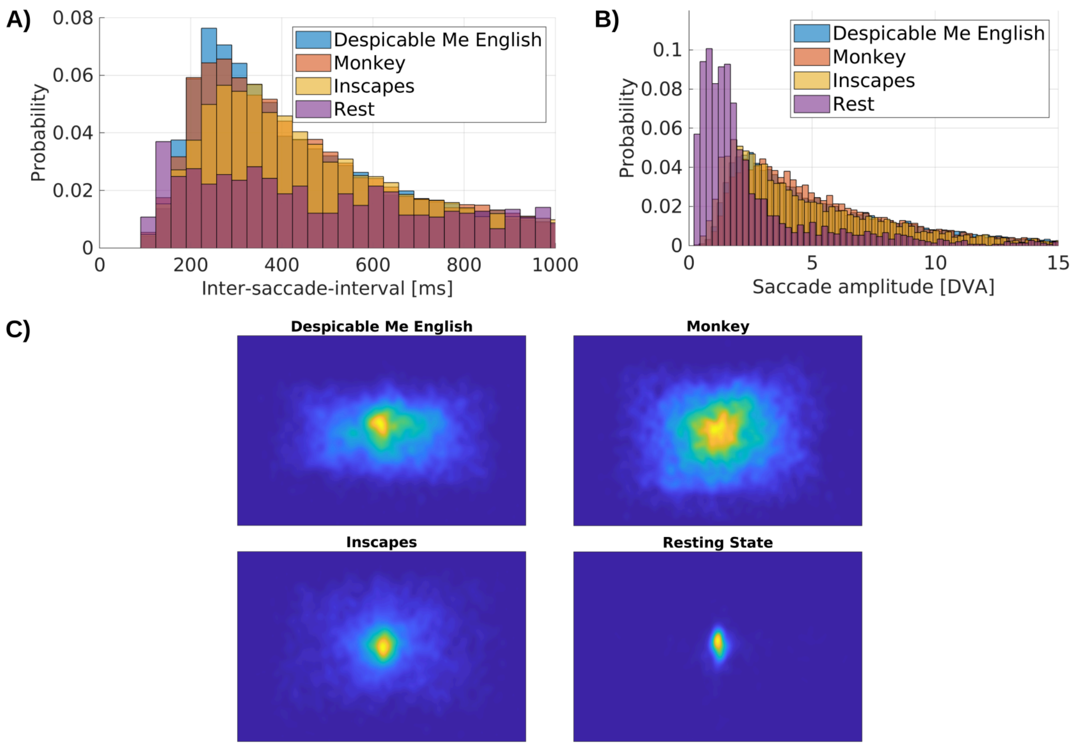


**Figure S6: Eye movement behavior differs between movies and resting state.** A) Normalized histogram of inter-saccade-interval, the time between consecutive saccades. B) Normalized histogram of saccade amplitude. C) Heatmaps of fixation position for different recordings. All figures are based on 10 min of data for 'Despicable Me English', 'Monkey' and 'Inscapes' and 5 min of Resting state. 'Despicable Me English': N=13643 saccades across 24 patients, with one recording each; 'Monkey': N=12790 saccades across 23 patients, with up to 2 recordings; 'Inscapes': N=8562 saccades across 20 patients, with one recording each; 'Rest': N=1510 saccades across 22 patients, with one recording each.

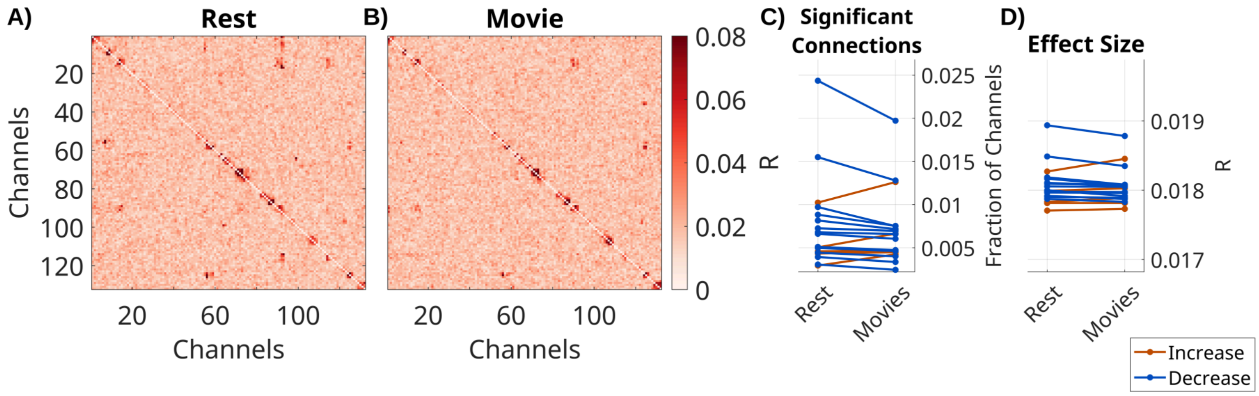


**Figure S7: Recurrent connectivity $\mathbf{A}$ of BHA decreases during movie watching compared to rest.** Same analysis as in Figure 4 with BHA data. Effect size $R$ for A) a VARX model during resting fixation with fixation onset as input feature. B) VARX model during movie watching, with sound envelope, acoustic edges, fixation onset and novelty, film cuts and motion as input features. C) Number of significant connections between movie and rest (fixed effect of stimulus: beta = $-5.4*10^{-4}$, t(88) = -2.1, p=0.042). D) Mean effect size across all channels between movie and rest (fixed effect of stimulus: beta = $-2.5*10^{-5}$, t(88) = -2.4, p=0.17). Each line represents one patient.

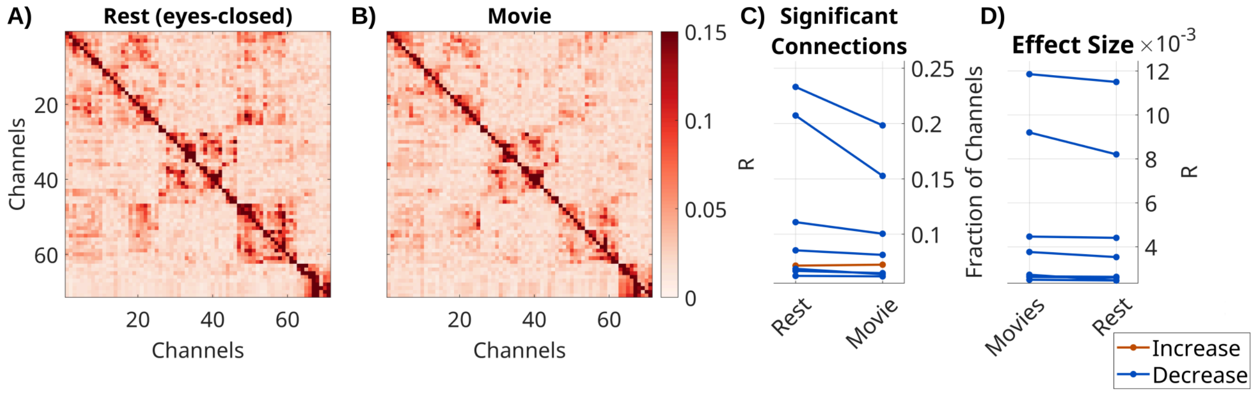


**Figure S8: Recurrent connectivity decreases in movies compared to eyes-closed rest.** Effect size $R$ for each connection in $\mathbf{A}$ in one patient (Pat_18_02) for A) a VARX model of 5 minutes of LFP recordings during movie watching, with sound envelope, acoustic edge, fixation onsets and novelty, motion and film cuts as input features. B) VARX model during eyes-closed rest without input features. Notably the majority of electrodes in this patient are located in the occipital cortex. C) The number of significant connections ($p<0.0001$) across all patients is lower during movie watching (median=-0.0017, p=0.01, N=8, Wilcoxon). D) Mean effect size across all channels is lower during movie watching (median=-0.0045, p=0.02, N=8, Wilcoxon). Each line represents one patient.

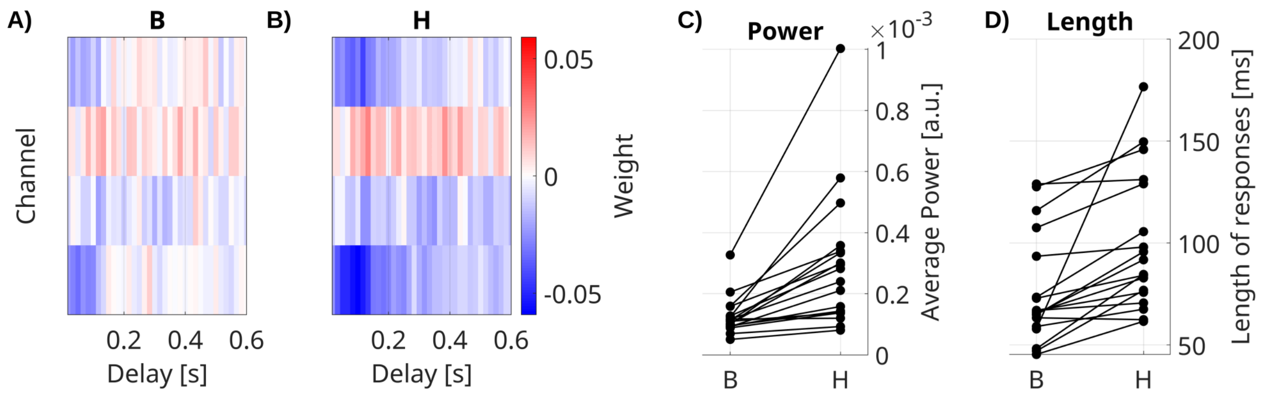


**Figure S9: BHA impulse response models.** A) Feed-forward BHA responses $\mathbf{B}$ to fixation onset are weaker and shorter than B) the overall system response $\mathbf{H}$. Significant responses for Pat_1. C) Power of responses $\mathbf{B}$ is weaker than $\mathbf{H}$ (medianΔ=-1.3*$10^{-4}$, p=0.0002, N=18, Wilcoxon). D) Mean length of responses $\mathbf{B}$ is shorter than $\mathbf{H}$ (medianΔ=-17.50ms, p=0.0002, N=18, Wilcoxon). Each line represents average values across significant channels in a patient.

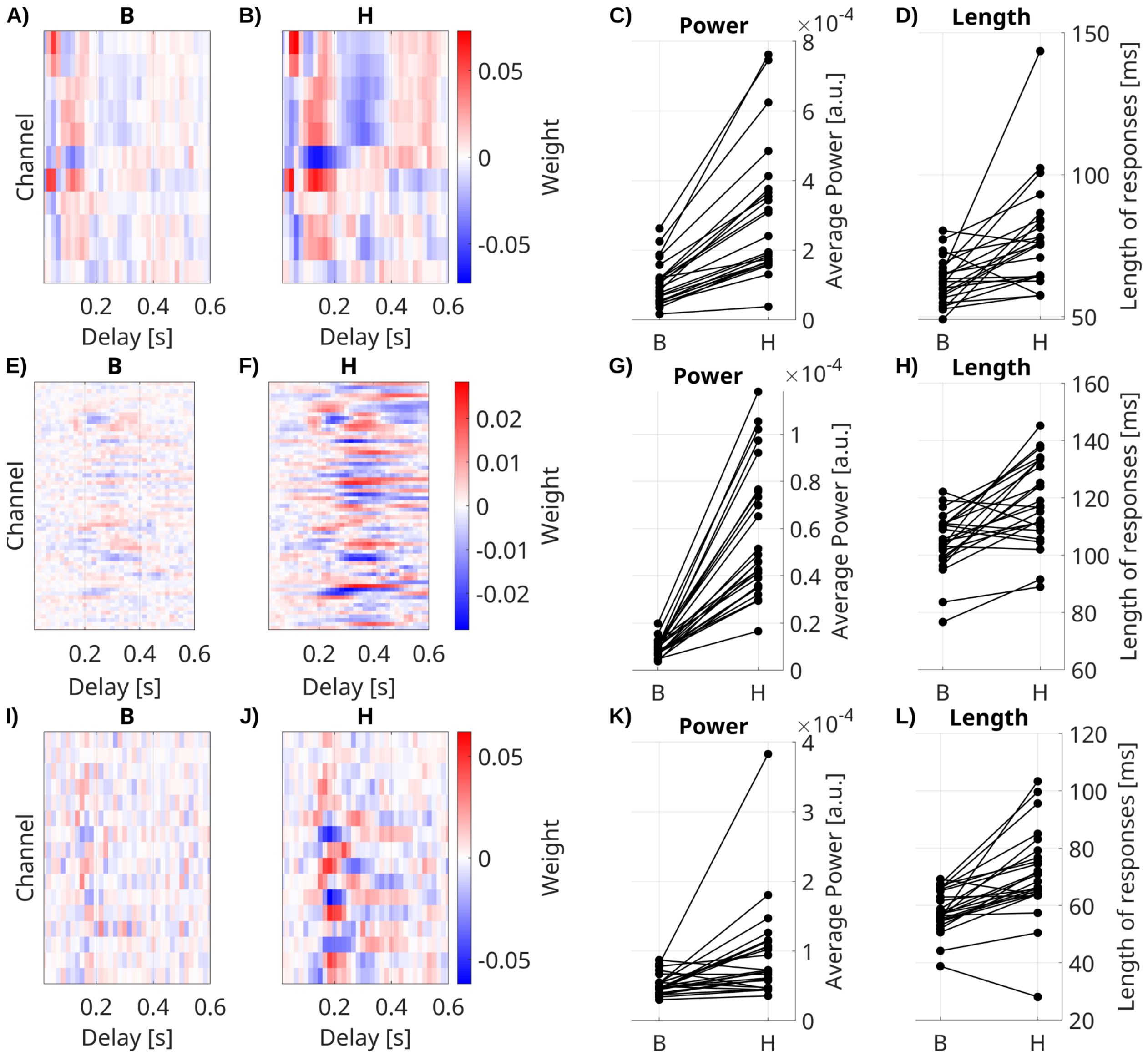


**Figure S10: Impulse response models.** Input responses to fixations onset (A-D), film cuts (E-H), and auditory envelope (I-L). Feed-forward responses $\mathbf{B}$ (A,E&I) are weaker and shorter than the overall system response $\mathbf{H}$ (B,F&J). Significant responses for Pat_1. Power of responses to fixation onset $\mathbf{B}$ is weaker than $\mathbf{H}$ for C) fixation onset (medianΔ=-1.5*$10^{-4}$, p<0.0001, N=23, Wilcoxon), G) film cuts (medianΔ=-3.9*$10^{-5}$, p=0.0001, N=25, Wilcoxon), and K) auditory envelope (medianΔ=-2.3*$10^{-5}$, p=0.0004, N=25, Wilcoxon). Mean length of responses to fixation onset $\mathbf{B}$ is shorter than $\mathbf{H}$ for D) fixation onset (medianΔ=-10.9ms, p<0.0001, N=23, Wilcoxon), H) film cuts (medianΔ=-14.98ms, p=0.0002, N=25, Wilcoxon), L) and auditory envelope (medianΔ=-10.53ms, p<0.0001, N=25, Wilcoxon). Each line represents average values across significant channels in a patient.

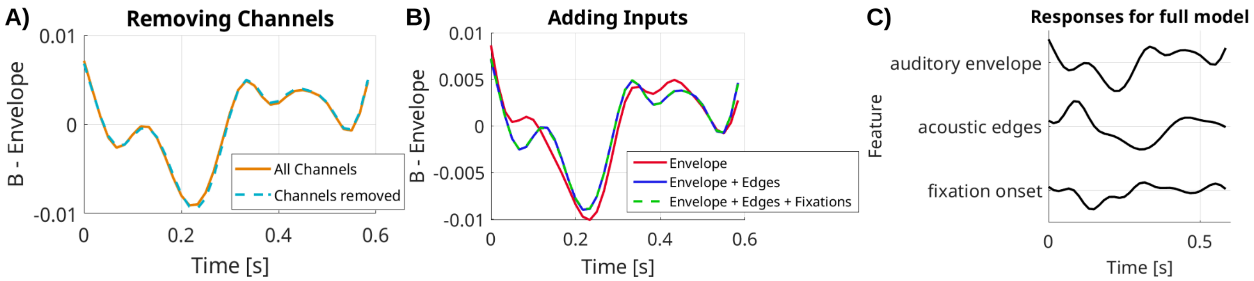


**Figure S11: Robustness of estimates of input filters $\mathrm{B}$ to intrinsic effects and uncorrelated features.** A) Removing half of all channels that do not show responses to the auditory envelope does not change the estimate of the response filter $\mathrm{B}$. B) The estimate of the responses $\mathrm{B}$ to the auditory envelope (red line), is changed by adding a correlated feature (acoustic edges, blue line), but not by adding an uncorrelated feature (fixation onset, green dashed line). C) The example channel shown in panels A and B, shows responses to the auditory envelope, acoustic edges and fixation onset. For a clearer visualization purposes responses have been filtered with a 10 Hz lowpass filter.

### Drop in LFP power for signal and innovation with sensory stimulation

We see a clear drop in power of the innovation e(t) over the entire spectrum analyzed for many channels (example of one patient in Fig. S12B). When looking at the median electrode we see that the effect is significant across all patients (Fig. S12C, Wilcoxon signed rank test p=0.005, N=25 patients). The drop in variability is evident also in the raw signals y(t) (Fig. S12C, Wilcoxon signed rank test p=0.011, N=25). It is most pronounced in oscillatory bands, for instance, in the particular patient shown in Fig. S12A there is a clear reduction in theta band activity around 8Hz. The specific bands differ across patients and channels (not shown). In total, even after oscillatory activity is modeled with the recursion filters A, there is a broadband reduction in power, while the relative noise power is unchanged (p>0.1).

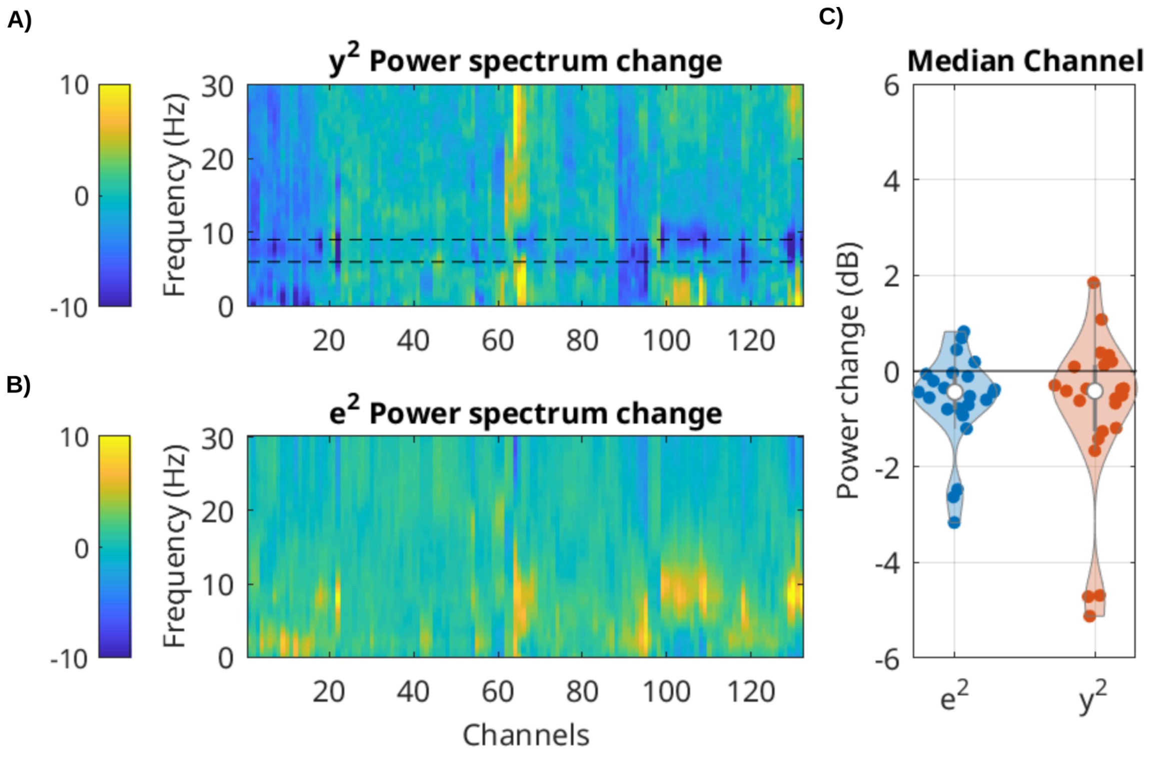


**Figure S12: For LFP, power of the signal and innovation process drops during movies as compared to rest.** Difference in LFP power between movies minus rest. Negative values indicate stronger power during rest. A) Difference in power spectrum for the raw signal y(t) for one patient. False color indicates change in the power spectrum in dB (blue hues indicate stronger power during rest). Dashed black lines bracket 5 and 11 Hz. B) Difference in power spectrum for the innovation process e(t). C) Power difference movie minus rest for each of 21 patients (each point is the median over channels).

### Grain adaptation model

This is a VARX model, where at every step the activity is adapted to have constant power over a given time horizon. We implemented this as a divisive normalization with by a running estimate of the power in the signal as follows:

$$\tilde{\mathbf{y}}(t) = \mathbf{A} * \mathbf{y}(t-1) + \mathbf{B} * \mathbf{x}(t) + \mathbf{e}(t)$$
$$|\mathbf{g}(t)|^2 = (1-\gamma)|\mathbf{g}(t-1)|^2 + \gamma\tilde{|}\mathbf{y}(t)|^2$$
$$\mathbf{y}(t) = \tilde{\mathbf{y}}(t)/\mathbf{g}(t)$$

The division with the gain $\mathbf{g}$ is element-wise. In the simulation here and in the main text we used $\gamma = 0.001$. This corresponds to power averaged over time with an exponential decay window with a time constant of $\tau = \Delta t/\gamma$, where $\delta t$ is the sampling interval. The simulation of Fig. S13 shows that a signal generated with this gain adaptation mechanism will exhibit the reduction of relative power of innovation (noise quenching) when the stimulus comes on, relative to when there is no external stimulus. But this is only true if the underlying signal generation implements gain adaptation (compare Fig. S13C vs S13D).

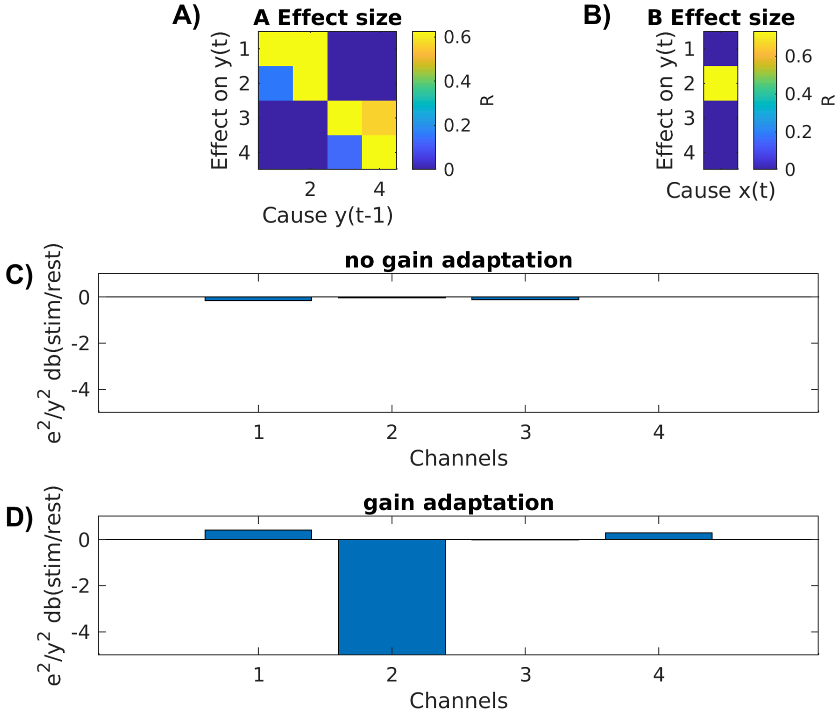


**Figure S13: Gain adaptation on a toy example.** In this small recurrent network there are 4 nodes, with each of 2 nodes connected. Input only arrives at one node. Data is simulated with and without gain adaptation, and then estimated with the VARX model. (A) Estimated recurrent connectivity (B) Estimated input connectivity estimated during “stimulus” condition. (C) Relative power of the innovation e (relative to signal y) subtracting dB between stimulus - rest condition. “Rest” here means that the input x was zero.

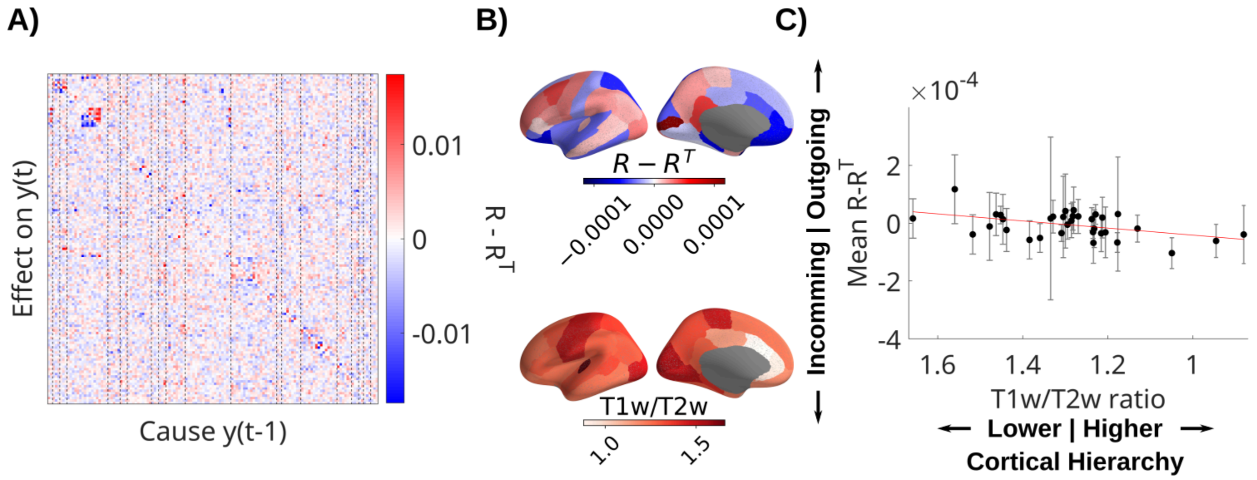


**Figure S14: Directionality of recurrent BHA connectivity in relation to cortical hierarchy.** Analysis as in Fig. 7. A) Difference of $R - R^\top$ showing asymmetric directed effects. Dashed lines indicate regions of interest in the Desikan-Killiany atlas. B) Mean directionality across patients and T1w/T2w ratio are averaged in parcels of the Desikan-Killiany atlas. C) Mean directionality is not significantly correlated with cortical hierarchy, estimated with the T1w/T2w ratio (t(533)=2.19, p = 0.029).
